# Lysine acetylation of PABPC1 C-terminal domain facilitates competitive recruitment of translation termination factor eRF3a

**DOI:** 10.64898/2026.01.10.698208

**Authors:** E Dubiez, YT Yeung, T Blee, M Soetens, S Virdee, AG Cook, NK Gray, M Brook

## Abstract

In eukaryotic cells, almost all mRNA transcripts are polyadenylated in the nucleus, with the poly(A)-tail being critical for their export, translation and stability. PABPC1 is a multifunctional RNA-binding protein (RBP) that binds poly(A)-tails and is a key driver of mRNA post-transcriptional regulation. It achieves this in part by interacting with a broad range of PAM2-motif containing proteins that bind the same common site within PABPC1’s PABC domain but result in diverse regulatory outcomes. Regulation of this intricate network of interactions remains poorly understood but our previous studies revealed the regulatory potential of mutually-exclusive acetylation or dimethylation of a key lysine sidechain (K606). To address this, we undertook a systematic study of PAM2-motifs, identifying translation termination factor eRF3a and translation repressor PAIP2 as having a >ten-fold higher affinity for PABPC1 relative to eleven other PAM2-motif partner proteins. Acetylation of PABPC1 at K606 specifically enhances its affinity for eRF3a *in vitro,* and we present the crystal structure of the specifically K606 acetylated C-terminal domain of PABPC1 in complex with the PAM2 motif of eRF3a, providing structural and functional insights into how acetylation facilitates competitive eRF3a recruitment to PABPC1 and mRNA. We further demonstrate that K606 acetylation promotes PABPC1-eRF3a interaction in cells, thereby antagonising other PAM2-dependent RNA effectors. Taken together, our results show that acetylation of lysine 606 in the PABC domain can specifically modulate PABPC1 partner selectivity, providing crucial insight into how PABPC1 multifunctionality can be regulated to coordinate mRNA usage and fate in the cytoplasm.

## Introduction

Post-transcriptional control of gene expression is a complex, multi-layered process that ensures precise cellular function^1^. RNA-binding proteins (RBPs), play a key role, influencing the fate of mRNAs from synthesis to degradation^2,3^. In eukaryotes, mRNA polymerase II transcripts normally exhibit a cap structure at the 5′ end and a 3′ poly(A) tail; both features that are important for post-transcriptional control and mRNA fate^4^. In the cytoplasm, these features stabilize mRNAs and enhance translation, putatively via mRNA 5′-3′ closed-loop formation. This involves interactions between the eIF4F translation initiation complex, associated with the mRNA 5’ cap structure, and the cytoplasmic poly(A)-binding protein 1 (PABPC1), bound to the poly(A)-tail^5^.

PABPC1 (PABP1) is an essential RBP in vertebrates, comprising four non-identical RNA recognition motifs (RRMs), an intrinsically disordered domain, and a PABPC1 C-terminal domain (PABC/MLLE)^6,7,8^. PABPC1 is the best studied member of the poly(A)-binding-protein family, which also includes tissue- and organism-specific isoforms. Initially described for its global roles in mRNA translation and stability, PABPC1 also acts as a scaffold protein that can recruit various effector proteins involved in different, and sometimes antagonistic, global and mRNA-specific aspects of mRNA fate. Many key effectors are recruited via a common peptide motif (PABP-interacting motif 2 or PAM2) that is recognised by the PABPC1 C-terminal domain (Figure 1A). PAM2 proteins can enhance (PAIP1)^9,10^ or repress (PAIP2) global translation initiation^11^, promote poly(A) tail removal (PAN3, TOB1/2^12,13,14^), stimulate translation termination and suppress nonsense-mediated decay (NMD) (eRF3a)^15,16^. PAM2 proteins can also positively or negatively control mRNA-specific translation (LARP1)^17^ and/or stability (ATXN2, TOB1)^18^ (Figure 1A). More than 10 human proteins containing PAM2 motifs (or variant PAM2) have been identified^19^, many are ubiquitously expressed, and all bind the same site within the highly conserved PABC domain^20^ likely competing for recruitment to PABPC1. This underscores the complexity of regulating PABPC1 function for mRNA fate decisions and it is unclear how the interplay between PABPC1 and PAM2-containing proteins is orchestrated^21^.

**Figure 1.**
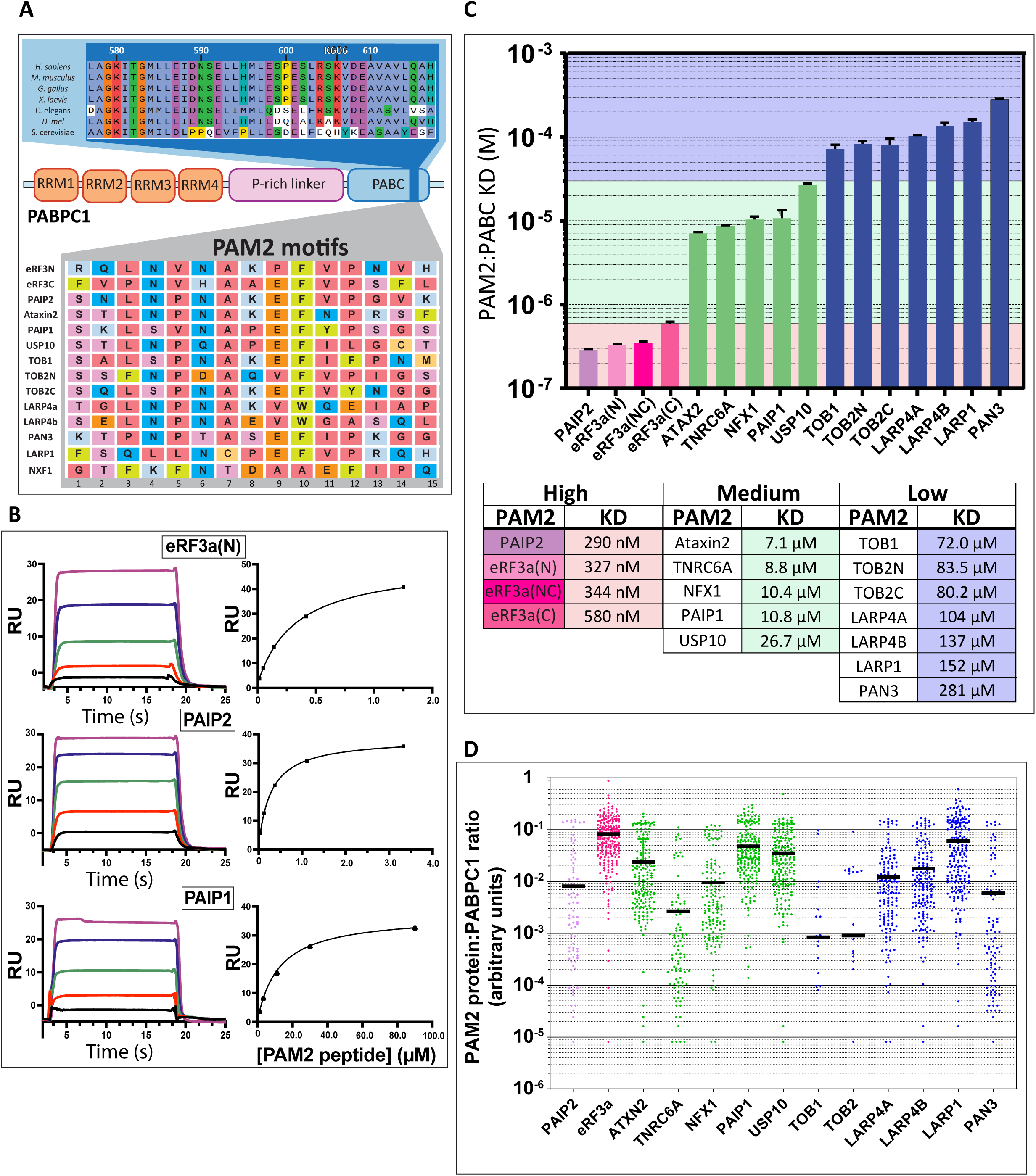
**(A)** PAM2 motifs from 12 different proteins all bind to the same region of the highly conserved PABPC1 PABC domain. **Upper**; blue panel depicts Hs PABPC1 amino acid sequence 577-617. Residues highlighted black are fully conserved between PABPC1in multiple species and between Hs PABPC2 and PABPC4. **Lower**; grey panel depicts PAM2 motifs using the L3 and F10 motifs of eRF3N as references. Residue colours: Red, aliphatic; Green, aromatic; Orange, acidic; light blue, basic; Pink, hydroxylic; light orange, sulfur-containing; dark blue, amidic. **(B)** Steady-state analyses of PAM2 motifs binding to PABC-His domain immobilised on NTA sensor chip by SPR. **Left;** Representative sensorgrams (shown as solid lines). Data was recorded for the association (20s) and dissociation (5s) phases for the various PAM2-motifs concentrations. **Right;** Representative diagram, A Hill model was used to fit the binding curve. **(C) Top;** Calculated K_D_ values from SPR analysis of the interaction of 16 PAM2 motif peptides, from 13 proteins, with PABPC1 PABC domain. **Bottom;** Table of mean calculated K_D_ values. Pink, ‘High’ affinity PAM2 motifs (K_D_ <6 x 10^-7^ M); Green, ‘Medium’ affinity PAM2 motifs (K_D_ <3 x 10^-5^ M); Blue, ‘Low’ affinity motifs (K_D_ <3 x 10^-4^ M). **(D)** PAM2 protein:PABPC1 expression stoichiometries. Calculated from matched PaxDB quantitative mass spectrometry data (downloaded 9/9/25). All PAM2 proteins are expressed at lower concentrations than PABPC1 in the same samples (see Supplementary Table 1 and Supplementary Figure 1).

A model was proposed for at least 3 PAM2 proteins, where interaction of the translation termination factor eRF3a with PABPC1 would antagonise the recruitment of PAN3 and TOB proteins, thereby protecting an mRNA against de-adenylation and decay, promoting translation^22^. However, such a model does not consider the other PAM2-motif proteins that can compete for PABPC1 interaction and the many other PABPC1 regulatory functions.

We previously identified multiple human PABPC1 post-translational modifications (PTMs), many of which are conserved from yeast^23^, and particularly noted multiple lysine residues that were subject to mutually exclusive acetylation or dimethylation. PTMs increase the number of proteoforms present in cells and, as they can be dynamic, expand the functional repertoire and regulation of proteins. Intriguingly, we identified PABPC1 lysine 606, which is a key PABC domain residue^23,24^ (Figure 1A), as being subject to acetylation or dimethylation and thereby being a potential site of regulation of PAM2 protein interaction.

In this study we systematically characterised PABC–PAM2 motif interactions using a unified experimental framework, with parallel methodological workflows across all assays. Our quantitative binding analyses reveal that among PAM2 motif–containing proteins, the translation termination factor eRF3a and the translational repressor PAIP2 exhibit the highest affinities for the C-terminal domain of PABPC1. Both PAM2s exhibit at least 10-fold stronger binding to PABC compared to those from 11 other PAM2 motif-containing interactors. Crucially, we further reveal that site-specific acetylation of a critical PABC lysine (K606) selectively strengthens PABPC1 interaction with eRF3a *in vitro*, whereas K606 dimethylation does not similarly impact PABC-PAM2 motif interactions. Furthermore, we report the crystal structure of the acetylated PABPC1 C-terminal domain bound to the eRF3a PAM2 motif. This provides high-resolution molecular detail on PABC surface remodelling that promotes competitive recruitment of eRF3a, thereby antagonizing other RNA-regulatory effectors. Finally, we detect substoichiometric levels of both K606 acetylation and dimethylation in cells and show that, as *in vitro*, eRF3a preferentially associates with acK606 PABPC1, indicating PTM-dependent regulation of PABPC1-PAM2 interactions *in cellulo*.

## RESULTS

### eRF3a and PAIP2 are the strongest PAM2 motif-mediated PABPC1 binding partners

The C-terminal PABC domain that interacts directly with PAM2 motifs^25^ is conserved from human to *S. cerevisiae*, reflecting its functional importance (Figure 1A). Many PAM2 proteins are ubiquitously expressed and known to bind directly to the same surface of PABPC1 and yet regulate different post-transcriptional processes. We hypothesized that differences in affinity of PAM2 motifs for PABPC1 would reveal a competitive hierarchy of PABPC1 interactions. Prior studies have only analysed a subset of interactions, used a variety of biophysical approaches and substrates, and reported a wide range of binding affinities, with K_D_ (dissociation equilibrium constant) values spanning from 0.2 to 40 μM^19,26^. To better understand the competitive interplay between the different PAM2 containing effectors, we applied a systematic approach, using Surface Plasmon Resonance (SPR) and identical experimental conditions to interrogate and compare binding affinity of 16 PAM2 peptides from 13 PAM2 proteins for the immobilised PABC domain (Figure 1B). Our study revealed binding constants (K_D_) spanning a 1000-fold range, from 300 nM to 300 μM allowing us to stratify the PAM2-containing proteins according to affinity (Figure 1C). Unexpectedly, only PAIP2 (290 nM) and the two eRF3a PAM2 motifs (eRF3a(N) and eRF3a(C)) (327 nM and 580 nM respectively) exhibited similarly high affinity for the PABC domain. In contrast, Ataxin2 (7.1 μM), TNRC6A (8.8 μM), NFX1 (10.4 μM), PAIP1 (10.8 μM) and USP10 (26.7 μM) displayed ∼20-80-fold lower K_D_ values (Figure 1C). The remaining PAM2 motifs exhibited very low μM affinity for the PABC domain: TOB1 (72.0 μM), TOB2N (83.5 μM) and TOB2C (80.2 μM), LARP4A (104 μM), LARP4B (137 μM), LARP1 (152 μM) and PAN3 (281 μM). These low affinity binders would be weak competitors for PABC when present at comparable concentrations to other PAM2 proteins.

These findings support the view that eRF3a and PAIP2 are likely the most competitive interactors for PABC domain binding. Notably, eRF3a harbours two overlapping, high-affinity PAM2 motifs (designated eRF3a(N) and eRF3a(C)), which enhance binding through avidity^27^. Once recruited to multimers of PABPC1 on a poly(A)-tail, PAM2 proteins are likely to undergo efficient rebinding to adjacent PABC domains, enhancing interactions in cells^28^. Such avidity-driven binding would confer a competitive advantage over other PAM2 proteins unless they are present at substantially higher local concentrations.

Analysis of quantitative proteomic datasets from a wide range of tissues and cell lines^29^ revealed that expression levels of all PAM2 proteins are ratiometrically orders of magnitude lower compared with PABPC1 (Figure 1D, Supplementary Table 1 and Supplementary Figure 1A) (ranging from ∼0.00084:1 (TOB1) to ∼0.082:1 (eRF3a) mean ratio). Principal component analysis of the sample datasets revealed similar PAM2 expression patterns across samples, i.e. there is no apparent subset clustering that would indicate divergent PAM2 protein expression patterns (Supplementary Figure 1B). Intriguingly, PAIP2 and eRF3a that have similarly high affinity PAM2 motifs, show ∼10-fold different mean ratio to PABPC1 (∼0.008:1 vs. ∼0.082:1, respectively, Figure 1D). Of note, eRF3a shows the highest mean ratio of expression to PABPC1 of all PAM2 proteins analysed (p <0.005 for all pairwise comparisons to other PAM2 proteins). These findings suggest that competition among PAM2-containing proteins for PABPC1 binding may be governed both by relative abundance, given their uniformly low cellular concentrations relative to PABPC1, as well as binding affinity. However, the presence of PTMs within the PABC domain raises the possibility that specific PAM2–PABPC1 interactions may also be dynamically regulated through PTMs that modulate PABC-PAM2 affinity or accessibility in response to signalling cues^30,31,32^.

### acK606 specifically increases affinity for eRF3a(N) PAM2 *in vitro*

We previously used mass spectrometry to identify multiple acetylated lysine residues in mouse and human PABPC1, and also mapped lysine, arginine, glutamate and aspartate methylation sites^23^; with many of these PTMs subsequently found in yeast^33^. Intriguingly, K606 which is critical for PABC-PAM2 interaction^34–36^, was detected in both acetylated (acK) or dimethylated (me2K) form, akin to histone methylation-acetylation ‘switches’. Based on molecular modelling we hypothesized that K606 PTM status could differentially alter PABC-PAM2 protein interactions^23^. Independent studies verified the existence of acK606-PABPC1^2,3^.

Characterisation of the role of K606 PTMs on PAM2 protein binding *in vitro* required methods to obtain PABC with a precise and quantitative modification at K606. We adapted an amber suppression approach, using an orthogonal pyrrolysyl-tRNA synthetase/tRNACUA pair to incorporate N-acetyl-L-lysine at the locus of K606^37^ (Supplementary Figure 2A). Unfortunately, this approach does not currently permit site-specific installation of dimethyl-lysine in recombinant protein. We therefore combined genetic code extension with a series of chemo-selective reactions to obtain specific K606 di-methylation within PABC^38^ (Supplementary Figure 2B). The mass of intermediary products was monitored by mass spectrometry (MS) (Supplementary Figure 2C).

The expected mass shift conferred by the PTMs was observed by MS (Supplementary Figure 2D) and the behaviours of modified recombinant proteins (acK606 and me2K606 PABC) were undistinguishable from the unmodified counterpart by size exclusion chromatograph (SEC), SEC-MALS and circular dichroism (Supplementary Figure 2E-H), indicating no unfolding, aggregation, or large structural changes. These results reveal that the PTMs (and all preparatory steps/processes) did not adversely affect the structure and monomeric nature of the proteins and that the samples were homogenously labelled.

To identify potential functional consequences of K606 acetylation or dimethylation on PABPC1-PAM2 interactions, we again used a systematic SPR strategy. To ensure unbiased quantification, all three modification states (unmodified, acK606, me2K606) of PABC were probed in parallel, using synthetic PAM2 peptides as analytes (Figure 2A, B). Me2K606 slightly increased PABC affinity for a subset of PAM2 peptides (TNRC6A, USP10, TOB2N, LARP1; all <1.2-fold) and slightly reduced binding to PAIP2 and LARP4B (>0.8-fold), whereas acK606 increased affinity for eRF3a, PAIP2, ATXN2, USP10, PAN3, and LARP4A and decreased affinity for TNRC6A PAM2 (Figure 2C). The fold-change decrease in TNRC6A binding to acK606-PABC was <0.7-fold, whereas the K_D_ values for increased acK606-PABC binding were ∼1.2-1.4-fold compared to unmodified PABC domain, except for eRF3a(N) which exhibited a 2.2-fold higher affinity (K_D_150.7nM vs. 327.2nM) (Figure 2C, D). Taken together, this makes eRF3a-acK606-PAM2 the highest affinity PAM2-PABC interaction described for a cellular PABPC1 protein partner. Given that eRF3a is expressed at the highest stoichiometric ratio with PABPC1, acetylation at K606 likely further increases its competitive advantage for PABPC1 PABC binding over other PAM2 proteins.

**Figure 2.**
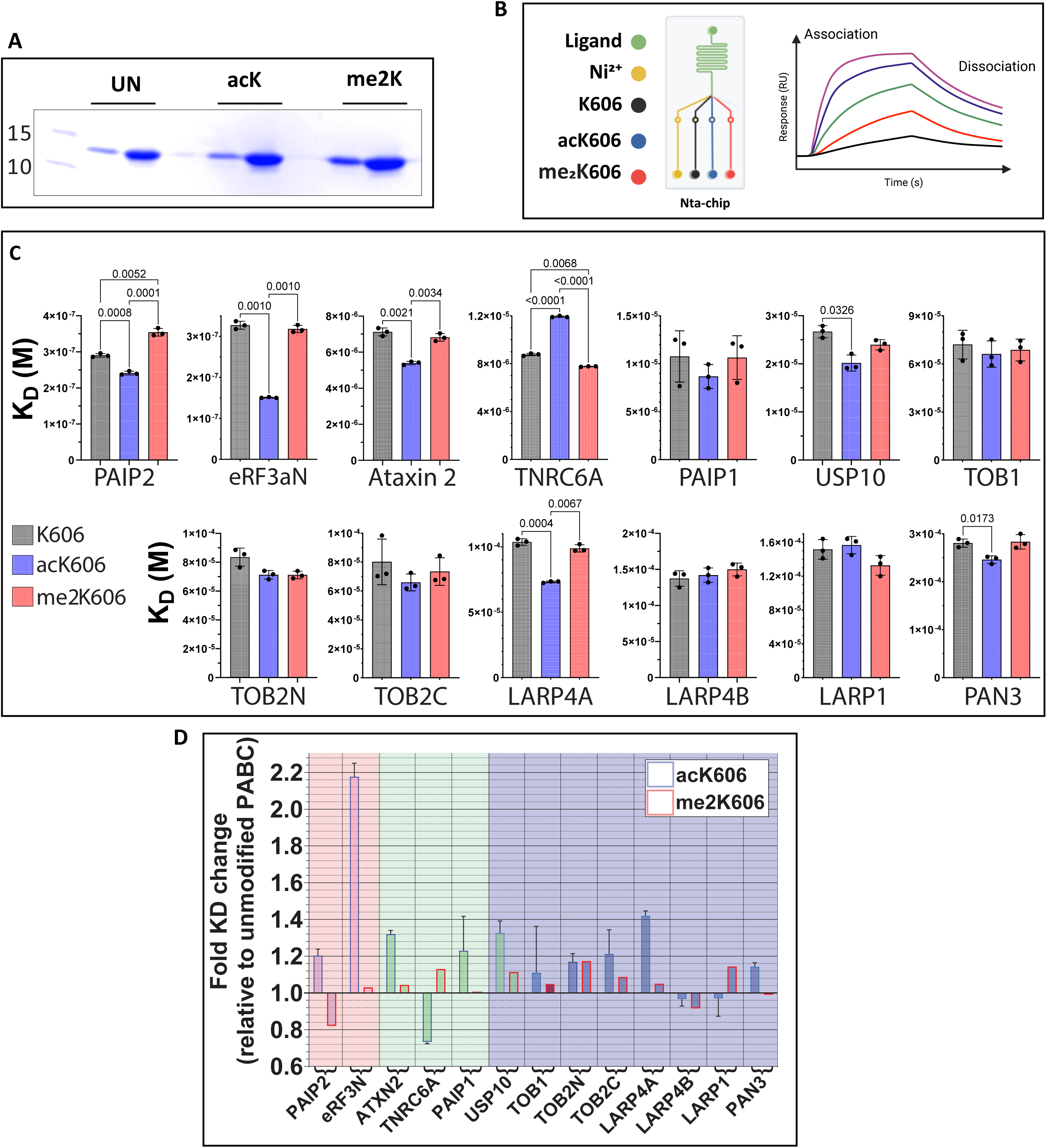
**(A)** SDS–PAGE analysis of purified recombinant PABC domains corresponding to unmodified, acetylated (acK606), and dimethylated (me2K606) forms. 1 and 5 µg of each variant was loaded on the gel. **(B)** Schematic representation of the SPR assay experimental setup used to simultaneously determine ligand affinities toward the different PABC variants. The NTA sensor chip was charged with Ni²□. One flow cell was left blank as a reference surface whereas the remaining three flow cells were immobilized with comparable amounts of each His-tagged PABC domain. Sensorgrams were recorded for the association (20 s) and dissociation (5 s) phases across a range of PAM2-motif concentrations (shown as solid lines). Equilibrium binding responses were fitted using a Hill equation to estimate apparent dissociation constants (K_D_). **(C)** Comparison of K_D_ for PAM2 motifs binding to PABC domains **(D)** Diagram representing K_D_ fold change relative to unmodified PABC for each PAM2 peptide. PAM2 motifs are in order consistent with Figure 1C: with Pink, ‘High’ affinity PAM2 motifs; Green, ‘Medium’ affinity PAM2 motif and Blue, ‘Low’ affinity motifs

### Crystal structure reveals molecular details of PAPBC1-acK606/eRF3a interplay

The structure of the unmodified PABC domain in complex with the human eRF3a(N) PAM2 motif was previously solved and refined to 2.3 Å^39^. To elucidate how PABPC1 K606 acetylation influences eRF3a(N) interaction, we determined the crystal structure of the acK606-PABC domain in complex with eRF3a(N). We obtained crystals in similar concentration, buffer conditions and space group symmetry as previously reported^39^. The crystal structure was solved by molecular replacement and refined to 1.8 Å (Supplementary Table 2). Comparison of the acetylated and unmodified structures revealed local specific rearrangement within the PAM2 binding region (Figure 3A, B). Acetylated Lys606 is stabilized by Glu592 via a hydrogen bond and electrostatic interactions, and is no longer coordinated with Glu609. Glu609 forms hydrogen bonds with Ser605 and water molecules. Overall, the structure of the acK606-PABC domain bound to eRF3a(N) is highly similar to that of the unmodified counterpart, with root mean square deviations below 0.902 Å across all atoms (Figure 3C, 3D).

**Figure 3.**
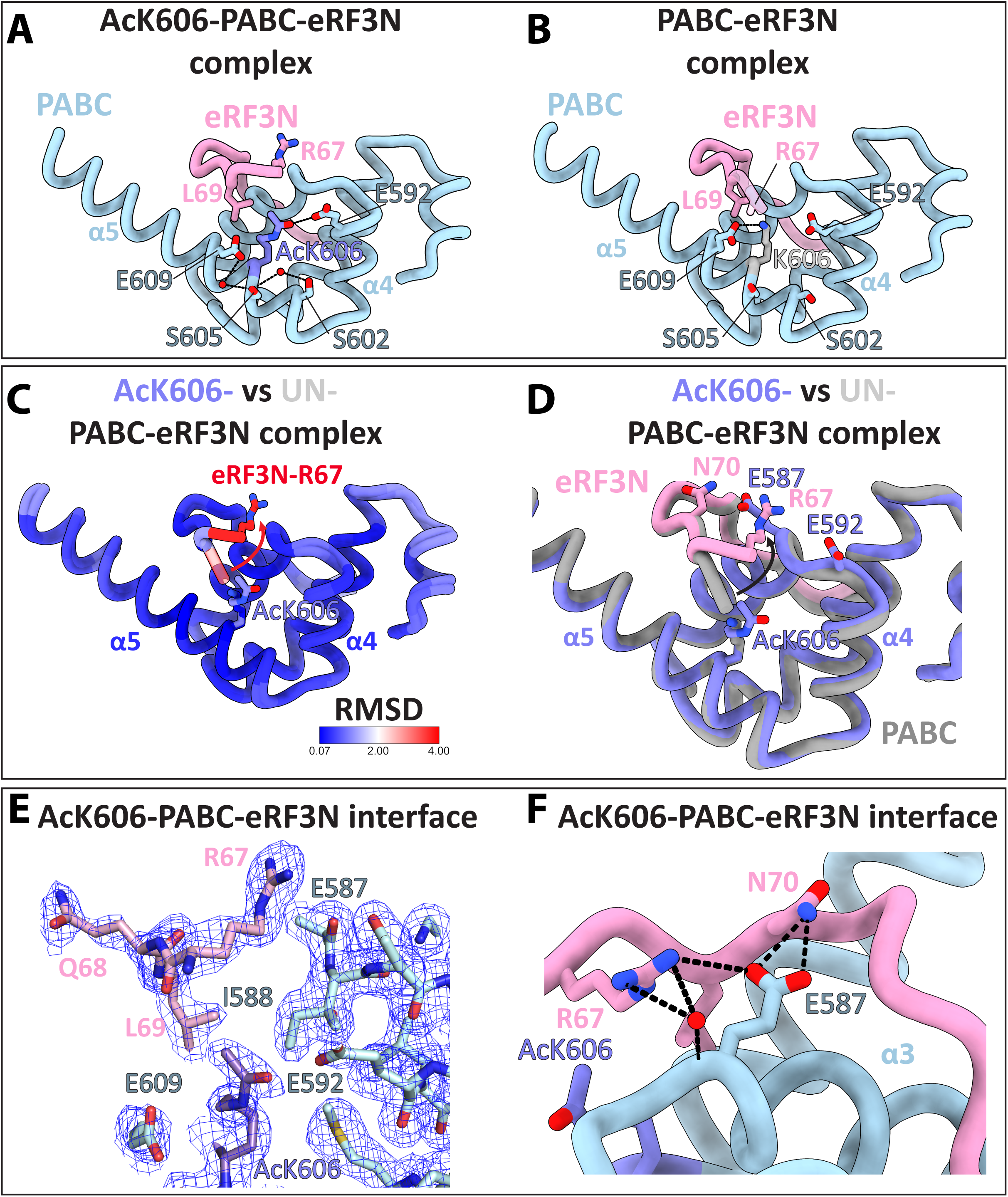
**(A)** Crystal structure of acK606-PABPC1–eRF3a(N), and **(B)** Crystal structure of unmodified PABC in complex with eRF3a(N) (PDB ID: 3KUI). PABC is coloured in light blue and eRF3 is depicted in pink, acK606 is in purple, Unmodified K606 is in light grey. Both structures are shown in the same orientation for direct comparison. Proteins are displayed as cartoon representations. PABC residues surrounding K606 are shown in blue sticks and labelled. Water molecules are represented as red spheres **(C)** Superimposition of the unmodified (PDB ID: 3KUI) and acK606-PABC-eRF3a(N) structures. Root mean square deviation (RMSD) with respect to the unmodified crystal structure is 0.902 across all atoms **(D)** Superimposition of the unmodified and acK606-PABC-eRF3a(N) structures. Colour coding follows that used in panel for the 606 residue: unmodified complex is shown in grey, acK606-PABC domain is shown in purple and its bound eRF3a(N) peptide is shown in pink, highlighting the conformational change of its N-terminal residue, depicted by a black arrow. **(E)** 2F□–F_c_ electron-density map contoured at 1.0 σ, illustrating the region surrounding the PABPC1–eRF3(N) interface at 1.8 Å resolution. The map was generated using the *carve* command in PyMOL. **(F)** Detailed view of the acK606-PABPC1–eRF3a(N) interface involving R67^eRF3a^ and N70^eRF3a^. The coordination network involving hydrogen-bonds or electrostatic interactions are indicated with dashed lines.

In the crystal structure of unmodified PABC-eRF3a(N) complex, eRF3(N) lies between alpha helices 4 and 5, anchored by binding of Leu69 ^eRF3a^ (Figure 3B). The side chains positions of Arg67-Gln68^eRF3a^ are poorly defined, indicating their high flexibility. In our structure of the acK606-PABC-eRF3a(N) complex, electron density for the side chains of Arg67-Gln68^eRF3a^ is clearly defined (Figure 3E). The position of Arg67^eRF3a^ lies between alpha helices 3 and 4 of PABC1 when the acK606 modification is present (Figure 3D, F). Arg67^eRF3a^ is stabilized by interactions with side chain and main chain atoms of Glu587^PABC^, and neighbouring water molecules (Figure 3F). Those contacts were previously absent in structures of the unmodified complex. Glu587^PABC^ also interacts with Asn70^eRF3a^ via hydrogen bonds. These additional contacts likely explain the increased affinity measured by SPR.

To better characterise acK606 mediated regulation of PABPC1-eRF3a binding, we recombinantly expressed full-length eRF3a^40^ (FL-eRF3a) (Figure 4A) and used a similar SPR approach to compare FL-eRF3a binding affinity towards unmodified/acK606/me2K606-PABC. Slower kinetics were observed using the full-length protein compared to peptides, which may reflect reduced flexibility of the PAM2 motif in the full-length context (Figure 1B, 4B). As with PAM2 peptides, K606 acetylation specifically increased the affinity of PABC domain for FL-eRF3a (2.4-fold), with K_D_ values of 1.3 μM (unmodified), 0.51 μM (acK606) and 1.49 μM (2meK606) (Figure 4C). Here, we observe that stronger FL-eRF3a interaction with acK606-PABC domain was potentially due to a faster binding rate during the ON phase (Figure 4D). To dissect the contribution of the different structural features observed in the acK606-PABC-eRF3a(N) structure, we assayed structure-based mutants of eRF3a (R67A and F85A) and investigated their interaction with unmodified/AcK/me2K-PABC domain by

**Figure 4.**
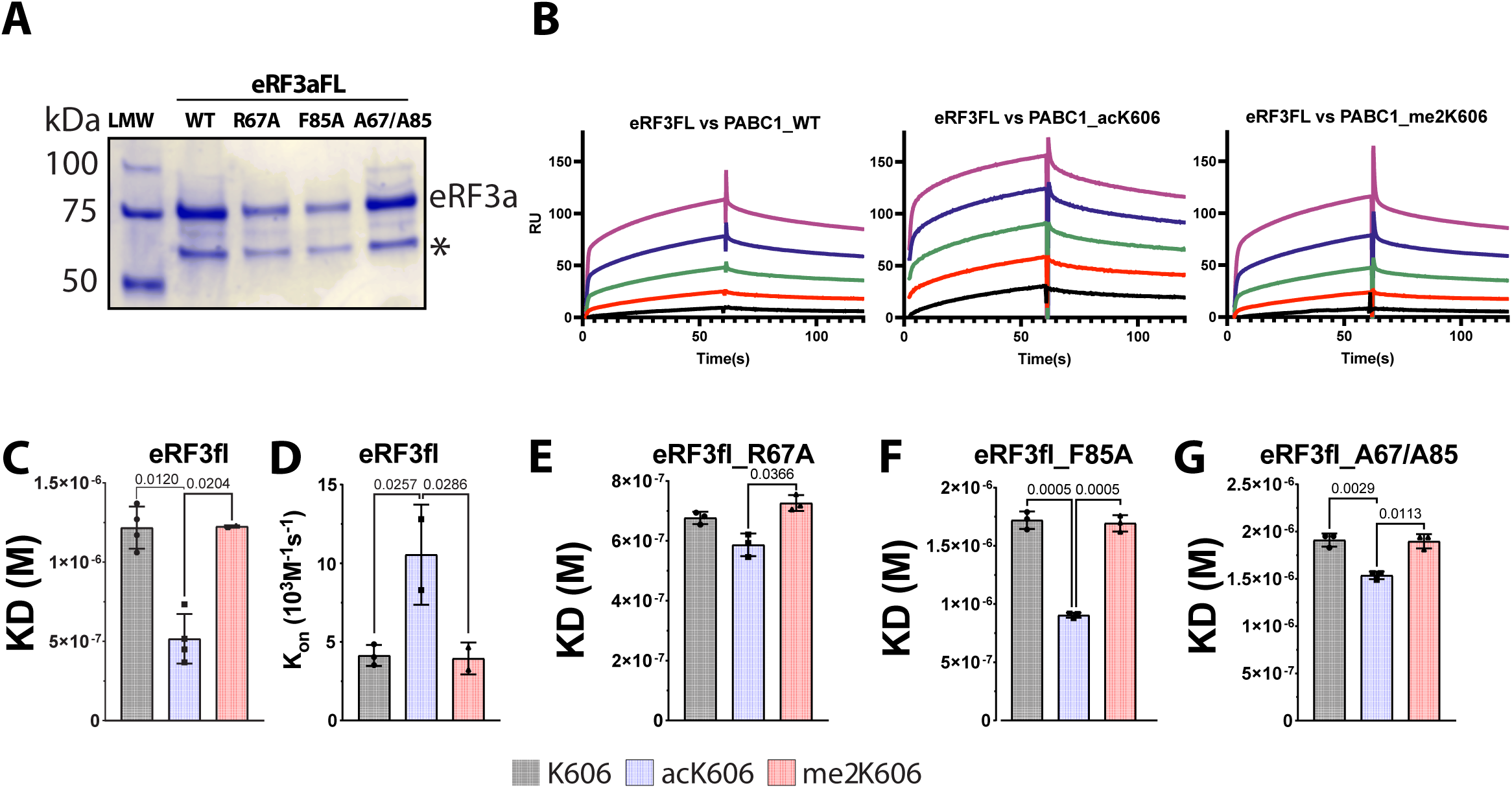
**(A)** SDS–PAGE analysis of purified recombinant full-length (FL) eRF3a wild type (WT) and mutants. * denotes previously seen contaminant^40^. **(B)** SPR analyses of FL eRF3a mutant binding to the His-tagged PABC domain. Representative experimental sensorgrams showing binding of FL eRF3a protein to unmodified and acK606 or me2K606 PABC domains **(C)** Apparent dissociation constants (K_D_) calculated as described previously are shown for FL eRF3a protein binding to PABC domains **(D)** Comparison of association rate constant (K_on_) for FL eRF3a protein binding to PABC domains. **(E-G)** Apparent dissociation constants (K_D_) are shown for each PABC domain with **(E)** eRF3a-R67A, **(F)** eRF3a-F85A, and **(G)** eRF3a-R67A/F85A double mutant.

SPR. eRF3a-R67A was selected because of the modification-specific interaction with acK606 PABC (Figure 3F). eRF3a is unique among PABPC1 partners in possessing two high-affinity PAM2 motifs (eRF3(N) and eRF3(C)) that bind overlapping sites on PABPC1. Despite this bivalency, eRF3a-PABPC1 complexes exist exclusively in 1:1 stoichiometry because both PAM2 motifs share a critical binding determinant: Phe76^eRF3a^, with only limited cooperativity between the two^35^. To selectively interrogate how acK606 modulates eRF3(N) binding in the context of full-length eRF3a, we introduced an F85A mutation which, based on prior evidence of this residue’s essential role in the eRF3a(C)-PABC interaction^35^, specifically disrupts eRF3(C) PAM2 binding. This separation-of-function mutation enabled us to isolate the contribution of the N-terminal PAM2 motif to acetylation-dependent binding. Additionally, we generated an R67A-F85A double mutant to evaluate the combined contribution of these residues.

All full-length mutant proteins could be successfully expressed and purified (Figure 4A). The R67A mutation abolished K606 acetylation-dependent binding, exhibiting comparable K_D_ values of 0.67 μM, 0.58 μM and 0.72 μM for unmodified, acK606 and me2K606 states, respectively (Figure 4E). This confirms that R67^eRF3a^ specifically mediates PABPC1 recognition in the context of K606 acetylation. By contrast, the eRF3a F85A mutant exhibited a binding profile similar to that of the WT-protein. eRF3a F85A K_D_ values range from 1.72 μM, 0.9 μM and 1.69 μM for unmodified-, acK606 and me2K606 states respectively, maintaining the acK606-specific affinity enhancement observed with eRF3a(N) PAM2 peptides (Figure 4F). The persistence of acetylation-dependent binding in the absence of a functional eRF3(C) PAM2 confirms eRF3(N) as the acetylation-sensing motif. Finally, introduction of the double eRF3a R67A-F85A mutations largely abolished the acetylation-dependant enhancement of eRF3a binding (K_D_ values of 1. 9 μM, 1.51 μM and 1.86 μM for unmodified-, acK606 and me2K606, respectively) (Figure 4G), as expected from the loss of the R67 side chain within eRF3(N). Wild-type eRF3a and the F85A mutant exhibit similar affinities for PABC variants, consistent with the limited cooperativity previously reported between the two eRF3a overlapping PAM2 binding sites^39^ and with the comparable SPR affinities measured for eRF3(N), eRF3(C) and eRF3(NC) PAM2 peptides (Figure 1C). Unexpectedly, we observed increased general affinity of eRF3-R67A mutant compared to WT, independently of the PTM state. We can speculate that while the arginine is critical for discriminating between modified and unmodified states, its presence may impose a conformational or electrostatic constraint that moderately limits overall binding compared to alanine mutant. This affinity enhancement must require an intact C-terminal interface, as it is absent in the R67A-F85A double mutant.

Collectively, this structure guided mutational analysis established R67^eRF3a^ as the acetylation-sensing residue. The crystal structure revealed how acK606 remodels the PABC PAM2-binding site, enabling additional molecular contacts between eRF3a(N) and a previously unrecognized interaction surface on PABC, anchoring the key determinant R67^eRF3a^ between helices 3 and 4 and thereby enhancing PABC-eRF3a binding.

### K606 acetylation favours cellular PABPC1-eRF3a interaction

To determine whether modification-dependent differences in PAM2-PABC affinity observed *in vitro* regulate cellular interactions, we substituted K606 to glutamine to mimic acetylation (K606Q)^41^ and also utilised two control variants: K606R (non-acetylatable unmodified mimic)^41^ and K606A (removes potential lysine side chain-specific interactions). Mimic residues (and their control mutations) can introduce unanticipated artefacts in protein structure and function, and we therefore performed extensive quality control on the recombinant mutant PABCs. The mutations did not alter recombinant PABC solubility or oligomerisation, as assessed by SEC coupled to multi-angle light scattering (Supplementary Figure 2F).

To validate the mimetic function of the K606Q mutation *in vitro*, and analyse K606A and K606R mutation effects, we introduced these substitutions in recombinant PABC domain and used SPR to compare the ability of the different K606X-PABC mutants to recruit eRF3a(N) PAM2 (Figure 5A). Neither the K606A nor the K606R mutations affected PABC binding affinities for eRF3aN, PAIP2, or PAIP1 PAM2 peptide, relative to PABC-K606 (native side chain), whereas the K606Q mutation effectively mimicked acK606 and caused an approximately 2-fold increase in binding affinity for eRF3aN only (Figure 5A). The small effect of acK606 on PAIP2 PAM2 binding was not replicated by the K606Q mutant. We then tested whether K606Q mutation could alter PABC affinity for full-length eRF3a and determined that PABC-K606Q exhibited a significantly greater increase in affinity relative to PABC-K606 than either K606A or K606R, both of which caused small increases (Figure 5A). Thus, K606Q-PABPC1 predominantly recapitulates the effect of acK606 and validates it as a mimic to study acK606 function *in cellulo*.

**Figure 5.**
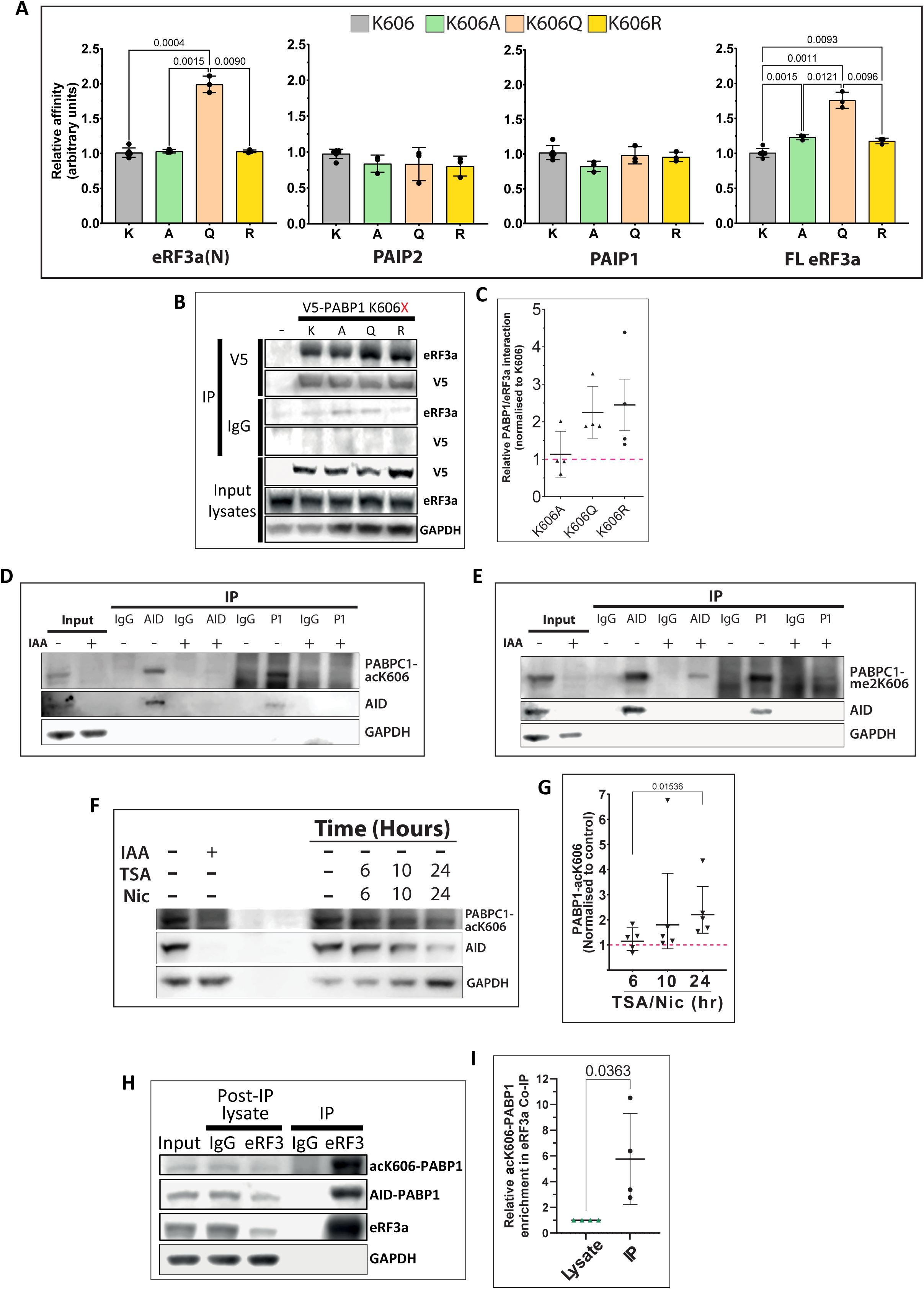
**(A)** SPR analyses of PAM2 motifs and full-length (FL) eRF3a protein binding to the PABC domain K606X variants. As before, His-tagged PABC variants were immobilised on an NTA sensor chip. Apparent dissociation constants (K_D_), calculated as described previously, are shown for each PABC variant. **(B)** eRF3a preferentially associates with acetylation-mimicking mutant K606Q-PABPC1 in cells. HCT-P1 cells were either untransfected or transfected with the V5-PABPC1-K606X mutants as shown. PABPC1 complexes were immunoprecipitated using anti-V5 antibody (or mouse IgG as negative control) and co-precipitating eRF3a was detected by immunoblotting. V5-PABPC1-K606X, eRF3a, and GAPDH levels are also detected in input lysates. **(C)** Dot plot depicting co-immunoprecipitated eRF3a levels relative to level in wild-type PABPC1-K606 IP (n = 4). Solid horizontal lines, mean; vertical solid lines, SD. **(D)** acK606-modified PABPC1 can be specifically detected. HCT-P1 cells were either left untreated (-) or treated with IAA for 2 hours (+) to induce PABPC1-AID degradation. PABPC1-AID was immunoprecipitated from cell lysates using antibodies against either the degron tag (AID) or PABPC1 (P1). Rabbit IgG was used as a negative control. Immunoprecipitates and input protein controls were immunoblotted for acK606-PABPC1, AID, and GAPDH. **(E)** me2K606-modified PABPC1 can be specifically detected. As (A) except immunoblotting is against me2K606-PABPC1. **(F)** PABPC1 K606 acetylation increases in the presence of deacetylase inhibitors. HCT-P1 cells were either left untreated (-), pre-treated with IAA for 2 hours (+) to induce PABPC1-AID degradation, or treated with 400nM trichostatin A/10mM nicotinamide for the indicated times. Lysates were immunoblotted for acK606-PABPC1, AID, and GAPDH. **(G)** Graph depicting PABPC1 K606 acetylation levels quantified relative to level of total PABPC1 at each time point and normalised to the ratio in untreated cells (n = 5). Solid horizontal lines, geometric mean; vertical solid lines, geometric SD; Kruskal-Wallis ANOVA analysis. **(H)** eRF3a preferentially associates with acK606-PABPC1 in cells. eRF3a-containing complexes were immunoprecipitated from HCT-P1 cells and co-immunoprecipitating total PABPC1-AID and acK606-PABPC1 levels were detected by immunoblotting. Input and post-IP lysates depict starting levels and depletion. Rabbit IgG was used as negative control. **(I)** Dot and whisker plot depicting acK606-PABPC1 levels quantified in input lysates and set to a value of 1 (Lysate). acK606-PABPC1 levels in eRF3a co-immunoprecipitates were normalised to the relative enrichment factor of total PABPC1 levels (IP). (n = 4) Solid horizontal lines, mean; vertical solid lines, SD (t test; *p* = 0.0363).

The same substitutions were introduced into PABPC1-V5 ectopically expressed in HCT-P1 cells, in which endogenous PABPC1 is tagged with an auxin-inducible degron (AID)^42^. Endogenous PABPC1-AID protein was depleted by addition of indole-3-acetic acid (IAA), enabling direct comparison of mutant PABPC1 properties *in cellulo*. PABPC1 relocalises to different sub-cellular compartments in response to a variety of cellular stresses but is predominantly cytoplasmic under steady-state conditions^43,44,45,46^.

We therefore assessed whether modification of K606 may alter PABPC1 subcellular distribution. Initially, we examined PABPC1 localisation by immunofluorescence in HCT-P1 cells. We found that endogenous PABPC1-AID and transiently transfected exogenous PABPC1-V5 were mainly cytoplasmic, indicating that neither AID nor V5 tagging altered PABPC1 steady-state localization^43^ (Supplementary Figure 3A). Consistent with effective auxin-induced degradation, PABPC1-AID was efficiently depleted upon IAA treatment. To test the effects of K606-PABPC1 modification-mimetic mutations, we transfected WT PABPC1-V5, K606A, K606Q, and K606R mutants and depleted endogenous PABPC1-AID to render ectopic V5-PABPC1 as the predominant PABPC1 in the cells. Immunofluorescence analysis revealed that exogenous WT and mutant PABPC1 exhibited similar predominantly cytoplasmic distribution, as seen for endogenous PABPC1 (Supplementary Figure 3B). Hence, based on these mimic analyses, modification of K606 is unlikely to alter bulk steady-state PABPC1 localisation.

We then examined the impact of K606 substitution on PABPC1–eRF3a association in cells using co-immunoprecipitation assays. V5-tagged wild-type PABPC1 or K606X variants were ectopically expressed as bait in HCT-P1 cells following depletion of endogenous PABPC1 (Figure 5B). K606A mutant co-purified similar amounts of eRF3a when normalized to V5-tagged WT PABPC1, whereas the acetylation-mimic K606Q mutant exhibited increased eRF3a binding, with K606Q consistently co-precipitating ∼2-fold greater amounts of eRF3a, thus validating its use as a mimic. The bulkier K606R mutant did not behave as expected and exhibited significant variability of interaction (Figure 5C) (see discussion).

To further investigate the potential role of K606 modifications in PAM2 protein binding, we raised antibodies against the K606 acetylated (acK606) and K606 dimethylated (me2K606) forms of PABPC1. These antibodies detected the two PTMs with high specificity in the site-specifically modified recombinant proteins (Supplementary Figure 2I). We then tested these antibodies in HCT-P1 cells and acK606-PABPC1 and me2K606-PABPC1 proteoforms were both detected in untreated HCT-P1 cells and became largely undetectable upon degron-dependent PABPC1 depletion (Figure 5D and 5E, Input lanes). The two proteoforms were also specifically enriched upon PABPC1 immunoprecipitation using two different antibodies (anti-AID and anti-PABPC1). K606 acetylation and dimethylation states are therefore both detectable in proliferating cells. To begin to evaluate whether acetylation of K606 could be regulated, HCT-P1 cells were treated with deacetylase inhibitors trichostatin A and nicotinamide for up to 24 hours (Figure 5F). acK606-PABPC1 was reproducibly detected in untreated cells. Prolonged deacetylase inhibition caused a reduction in overall PABPC1 levels as observed by loss of signal with anti-AID tag. Despite this, the average acK606-PABPC1:PABPC1 ratio increased upon treatment with deacetylase inhibitors treatment (Figure 5G). Taken together, these data indicate that PABPC1 can be acetylated or dimethylated on K606 *in cellulo*, and that steady-state acK606-PABPC1 levels are substoichiometric.

To directly assess the contribution of K606 acetylation to endogenous PABPC1–eRF3a association *in cellulo*, we performed co-immunoprecipitation experiments using endogenous eRF3a as bait (Figure 5H). This revealed a mean >5-fold enrichment of endogenous acK606-PABPC1 level in the eRF3a-bound fraction relative to the proportion of acK606 detected in total cellular PABPC (Figure 5I). This result indicates that PABPC1 K606 acetylation increases its interaction with eRF3a, even in a competitive cellular environment. Collectively, these findings establish that acetylation of PAPBC1-K606 favours the recruitment of the translation termination factor eRF3a in cells, in line with our *in vitro* and structural studies.

## Discussion

Our comprehensive analysis of PAM2 motif binding to PABPC1 *in vitro* revealed that, among the 13 human PAM2 motifs tested, only eRF3a and PAIP2 motifs bind with sub-micromolar affinity, from 20- to a 1000-fold tighter than any other PAM2 peptide assayed. Our data suggest a hierarchical recruitment model based on affinity, rather than a broad competition between equivalent PAM2 proteins. In this model, eRF3a and PAIP2 dominate recruitment to PABC under basal conditions. Furthermore, our analysis of quantitative proteomics data highlights large differences in relative ratios of PAM2 proteins to PABPC1, under physiological conditions in multiple cell/tissue types. PAM2-containing proteins are expressed at substantially lower levels than PABPC1 across diverse tissues and cell lines (ranging from ∼0.00084:1 to ∼0.082:1), suggesting that relative abundance may contribute to regulating competition for PABPC1 binding. For example, given their low abundance and their weaker affinity for PABC, it would be expected that PAM2 proteins like PAN3, TNRC6A, or TOB1/2 would effectively be outcompeted under normal conditions by eRF3a and PAIP2 and likely require one or more of increased local concentration effects, contributions from other partner proteins, or PTMs to facilitate interactions with PABPC1.

However, it is important to note that, for a small subset of PAM2 proteins (e.g.: PAIP1^47^, PAIP2^48,49^, LARP4^50^), the PAM2 motif is not the sole interaction interface with PABPC1, some may participate in complexes with other PABPC1-binding partners, and some PAM2 proteins also bind RNA. In particular, recent work by Sagae et al. demonstrated that PAIP2A inhibits translation by directly competing with RNA for binding to the RNA recognition motifs (RRMs) of PABPC1, thereby promoting dissociation of PABPC1 from poly(A) RNA^49^. Although that study focused primarily on regulation at the RNA-binding interface, it is notable that PAIP2 dissociated full-length PABPC1 from poly(A) RNA at least 10-fold more efficiently than a shorter construct that lacked the PABC domain, suggesting that PAM2–PABC interactions significantly contribute to the efficiency of PABPC1 displacement. These observations support a model in which PABPC1 activity is governed by combinatorial regulation across distinct domains, with RNA occupancy controlled at the RRMs and partner selection tuned at the PABC domain. Such multi-layered regulation provides a framework for how post-translational modification and competitive binding events are integrated to switch PABPC1 between translationally active and repressive states. Together, these results highlight potential niches for context□ or cell type-dependent and/or PTM□driven engagement, offering a mechanistic basis for stress and developmentally regulated shifts in mRNP composition.

Indeed, an alternative, non-mutually exclusive possibility is that specific PAM2–PABPC1 interactions are dynamically regulated through post-translational modifications (PTMs) of either the PAM2 motif and/or the PABC binding domain, modulating their affinity or accessibility in response to signalling cues^30,31,32^. Hence, we then focused on a previously under-appreciated layer of regulation within the PABPC1–PAM2 interaction network, the acetylation or dimethylation at Lys606 of PABPC1 and hypothesized that it could act as a molecular switch to direct effector selection on mRNA poly(A) tails^23^. While leaving interactions with other PAM2 factors unaffected or only minimally affected, this acetylation specifically increased the affinity of PABC for eRF3a, potentially due to a faster association rate. Faster binding in presence of acK606 might significantly enhance eRF3a’s ability to outcompete other PAM2 proteins, rapidly occupying and rebinding available binding sites on poly(A)-associated PABPC1s, leading to more stable complex formation. This kinetic advantage could reduce the likelihood of transient competitors forming stable interactions. This is especially true considering the presence of the second overlapping eRF3a(C) PAM2 motif^35^. The probability of complete dissociation decreases, and the avidity effect can shift the equilibrium towards the specific bound state. This is crucial in signalling pathways or regulatory networks where rapid and selective binding dictates downstream responses^35^.

We gained molecular insight into the mechanism of acetylation-enhanced PABPC1-eRF3a affinity from the crystal structure of the acK606-PABC domain in complex with the eRF3a PAM2 peptide. This revealed that acetylation induces a local rearrangement of acidic and hydrophobic residues, reshaping the eRF3a binding surface and favouring an interaction between R67^eRF3a^ and E587^PABC^^1^ that was not observed in an equivalent structure with unmodified PABC. Structure-guided mutagenesis of R67 supports its role in the acetylation-enhanced interface: its specific mutation abolished the effect of acK606 on affinity while a mutation affecting eRF3a C-terminal PAM2 binding did not. While we did not identify a specific function for K606 me2K, we speculate that it could block acetylation or other PTMs like ubiquitination.

From a functional perspective, we detected the substoichiometric (and likely reversible) acetylation or dimethylation in human cells and conducted endogenous co-immunoprecipitation experiments in which we observed a five-fold enrichment acK606-PABPC1 in association with eRF3a, reinforcing the physiological relevance of our *in vitro* findings. This increased eRF3a interaction was recapitulated by acetylation-mimicking PABPC1 (K606Q), providing a useful tool for future studies of this PTM. Interestingly, the K606R substitution exhibited highly variable behaviour. This observation reinforces the notion that lysine-to-arginine (K→R) substitutions are not strict mimics of unmodified lysine, as arginine, despite sharing a positive charge, has a bulkier side chain with distinct hydrogen-bonding geometry and capacity that can perturb protein–protein interactions at the interface. Such limitations of K→R substitutions have been noted previously in studies of PTM mimetics ^51,52^. It is also possible that PABPC1 K606R mutation is affecting interaction with other partners and artefactually altering the ability of eRF3a to compete for binding.

The affinity advantage imparted by acK606 likely has significant biological consequences. In cellular contexts where numerous PAM2 competitors coexist, modification-triggered recruitment of eRF3a may ensure efficient coupling of translation termination with mRNA protection against deadenylation. PAIP2 protein turnover is dependent on its interaction with PABPC1, which antagonises PAIP2 recruitment to the ubiquitin ligase EDD that targets it for degradation^54^. Stronger eRF3a-PABPC1 could not only enhance translation by antagonising the recruitment of the translation repressor PAIP2, but could consequently increase EDD-dependent PAIP2 degradation to either maintain or reinforce the stoichiometric advantage of eRF3a for PABPC1 binding. Furthermore, it was previously demonstrated that eRF3a antagonizes the deadenylase-recruiting factors Pan3 and Tob^54^. Acetylation at K606 may represent a regulatory node by which cells fine-tune mRNA stability and translational capacity under stress or developmental cues.

PABPC1 is commonly degraded by viral proteases to repress translation of endogenous proteins, but it can also be subjected to viral hijacking to promote translation of viral proteins. Mimicry of host small linear motifs (like PAM2) is ubiquitous viral strategy^55^. Recently, a study presented an engineered, “super” chimeric sPAM2 motif, using phage display combining different PAM2 factors domains. After screening up to 10^13^ potential peptides, sPAM2 exhibited a 10-fold improved affinity compared to best known PAM2 binder^57^ (close to K606 acetylation outcome). Our study reveals the importance of residues spanning outside of the classically described PAM2 peptides (e.g. eRF3a R67 to N70), in presence of PTM(s). Hence, it could serve as a basis to engineer even stronger, longer PAM2, for advancing novel peptide inhibitors to combat viral infections.

Beyond the immediate mechanistic insights, our work broadens the scope of post-translational control in RNA biology. Lysine acetylation and dimethylation have been extensively studied in the context of histone regulation, yet, despite their identification on a growing number of RBPs, they remain underexplored, likely due to the often very low stoichiometry of modification^58^ and the lack of highly specific tools for in cellulo studies. The weaker effects of acetylation and/or dimethylation on affinity of some PAM2 protein for PABPC1 may indicate further regulatory complexity remaining to be explored. However, the very low affinity of many of the affected PAM2s suggests that additional inputs would be needed to make these affinity changes have biological relevance. The discovery that PABPC1 appears to employ an acetylation-driven switch to manage competitive binding events suggests that similar PTM “codes” may operate across the landscape of RBPs, many of which are also multifunctional, to coordinate complex post-transcriptional networks. Future studies should survey the PABPC1 interactome under varying cellular states to identify upstream “writer” and “eraser” enzymes for K606 modification and other regulated partners, as well as downstream functional effects on global translation and mRNA turnover.

## Materials and methods

### Expression plasmids

Full-length human eRF3a (eRF3aFL) was subcloned into the pETM-20 expression vector (a gift from Günter Stier, EMBL Heidelberg) as previously described^40^. Briefly, the eRF3a coding sequence was amplified by PCR using primers containing NcoI and XhoI restriction sites and ligated into pETM-20 digested with the corresponding endonucleases, yielding to construct encoding eRF3a flanked by an N-terminal thioredoxin protein followed by 6His for affinity purification and a TEV cleavage site. To generate pET52b-Strep-PABC[544–636]-His expression constructs, the PABC coding sequence was amplified from full-length human PABPC1 using PABC-specific primers introducing KpnI (5′) and SacI (3′) restriction sites. The resulting fragment was cloned into the pET52b(+) vector, yielding constructs encoding the PABC domain flanked by an N-terminal Strep-tag II and a C-terminal deca-histidine tag. Full-length human PABPC1 WT and variants were cloned into mammalian expression vector pCI-MS2V5 (Addgene #65807) digested with XhoI and NotI enzymes using similar strategy. Point mutations in PABC and eRF3a variants were introduced by PCR-based site-directed mutagenesis using the QuikChange™ protocol (Agilent), following the manufacturer’s instructions.

### Protein expression and purification

BL21 (DE3) *E. coli* were transformed with pET52b-PABC plasmid and induced with 1mM Isopropyl β-d-1-thiogalactopyranoside (IPTG) at an OD_600_ of 0.7. Bacteria were grown overnight at 21°C before being harvested. Pelleted cells were resuspended in lysis buffer containing 20mM HEPES pH 7.0, 100mM NaCl, 2mM Dithiothreitol (DTT) complemented with Ethylenediaminetetraacetic acid-free (EDTA-free) proteinase inhibitor cocktail (Roche), 20mM imidazole, 10μg/ml RNase A and 10μg/ml DNase I were added to lysate. PABC-His was bound to Ni-NTA resin (Qiagen), washed with buffer containing 20mM imidazole and eluted with 500mM imidazole. Eluted PABC-His was run through a Resource Q FF column (Cytiva) using a linear NaCl gradient (100mM to 1M), before being applied to a Superdex 200 Hiload 16/60 size exclusion chromatography column (Cytiva) in GF buffer: 20mM HEPES pH 7.0, 150mM NaCl, 2mM DTT.

Full-length (FL) human eRF3a was prepared as previously described^40^. Briefly, BL21 cells were transformed with pETM-20-eRF3aFL plasmid and grown in 2xLB, supplemented with 100mg/ml carbenicillin at 37°C to OD_600_ 0.6. Cells were transferred to ice for 30 min and induced with 0.25mM IPTG at 20°C overnight. Cells were resuspended in lysis buffer 20mM HEPES pH 7.0, 100mM NaCl, 2mM DTT, 10% glycerol supplemented with 10μg/ml RNase A and 10μg/ml DNase I and 20mM imidazole. Cleared lysate was loaded onto a Ni-NTA resin (Qiagen) and washed in buffer containing 3M NaCl and eluted in buffer containing 100mM NaCl and 200mM imidazole. Eluted protein was further purified on a Resource Q FF column (Cytiva) using a linear gradient (0.1–0.8M NaCl). The eRF3a peak was collected, TEV protease added in 1:25 ratio, and dialyzed overnight against lysis buffer containing 0.1M NaCl. The TEV protease and the cleaved Trx tag were removed on Ni-NTA resin (Qiagen). Cleaved protein was further purified on Superdex 200 Hiload 16/60 size-exclusion columns (Cytiva).

### Protein Modification

For the generation of site-specifically acetylated and dimethylated PABC, pET52b-PABC vector containing an amber codon at position 606 was co-transformed into *E. coli* BL21(*DE3*) cells with pBK-AcKRS-pylT (expressing the orthologous tRNA synthetase) and pCDF-tRNACUA-pyIT (expressing the orthologous tRNA-CUA to recode the Amber STOP codon). Cells were grown at 37°C in LB medium containing 100µg/ml carbenicillin, 50µg/ml spectinomycin and 34µg/ml kanamycin. At OD_600_□=□0.4, LB was complemented with either 10mM acetyl-lysine (Merck) and 20mM nicotinamide (Merck) or 2.5mM Boc-Lys-OH. At to OD_600_□=□0.8, 1mM IPTG was added for overnight induction at 20°C. Modified PABC proteins were purified as per unmodified PABC.

To obtain me2K606-PABC, purified PABC-K606Boc was lyophilized. Protein was dissolved in 6M Guanidine-Cl (Gdn-Cl) and diluted with DMSO. To globally protect PABC’s 6 lysines and N-terminal amine, excess of Cbz-Osu (for N-(benzyloxycarbonyl)succinimide) (9:1) and N,N-diisopropylethylamine (DIEA) (90:1) were added and incubated with stirring for 1 hr at 23 °C. The protected polypeptide was then precipitated with ice cold ether (1:10 dilution). The precipitate was collected by centrifugation, washed twice with cold ether and air-dried. To specifically deprotect the encoded protecting Boc group, the globally protected protein was dissolved in cold 60% TFA/H_2_O and incubated at 4°C for 1hr. The PABC-(Cbz) protected protein was precipitated and washed with ice-cold ether. The aqueous and ether layers were removed and precipitate was air-dried. To specifically di-methylate K606, the protein was dissolved in 6M Gdn-Cl, 100mM phosphate buffer (pH 7.5) using sonication. Dimethylation of PABC was carried out using a JBS methylation kit (Jena Bioscience). After 4hr reaction, the K606 dimethylated protein was precipitated with ice cold ether, collected by centrifugation, pellet was washed twice with cold ether and air-dried. To deprotect cbz-protected lysine residues and N-terminal amine, protein was then dissolved in ice-cold 7:11:2 DMS/TFA/TfOH (^v^/_v_) and stirred for 2hr on ice. The protein was further purified by RP-HPLC using a gradient of 5%–75% Buffer B over 30 min at a flow rate of 5 ml/min. Buffer A = 0.1%TFA in H_2_O, Buffer B = 10% Buffer A in acetonitrile to obtain PABC-me2K606. The protein was precipitated with ice cold ether, collected by centrifugation and washed twice with ice cold ether. The dried precipitate was dissolved in unfolding buffer (8M urea, 100mM Na_2_HPO_4_ [pH 7.4], 500mM NaCl), dialyzed in the same buffer overnight at room temperature The protein was refolded via dialysis in purification buffer (20mM HEPES pH 7.0, 150mM NaCl, 2mM DTT) and refolded protein was purified on Superdex 200 Hiload 16/60 size-exclusion columns (Cytiva) for further use in biochemical assays (∼10% recovery yield).

### Crystal structure

Crystals of acK606-PABC domain bound to eRF3a were obtained by equilibrating a 1μl drop of an equimolar mixture of PABC with eRF3a peptide (10 mg/ml) in buffer (10mM MES and 100mM NaCl, pH 6.3), mixed with 1μl of reservoir solution containing 2.3M ammonium sulphate, 0.2 M sodium sulfate and 0.1 M sodium acetate at pH 5.4. Crystals grew in 2-5 days at 20°C. The solution for cryoprotection contained the reservoir solution with addition of 20% (^v^/_v_) glycerol. For data collection, the crystals were harvested in a nylon loop and flash cooled in a N_2_ cold stream. Data were collected at Diamond Light Source (DLS) on beamline I04-1. Data from a crystal diffracting to 1.8 Å, with space group *P* 2_1_ 2_1_ 2, were obtained and indexed and reduced using the automated data processing suite at DLS. The crystal contains one PABC and one PAM2 molecule per asymmetric unit. The structure was solved by molecular replacement, using the structure of unmodified PABC-eRF3N [Protein Data Bank (PDB) ID: 3KUI], followed by rounds of rebuilding in COOT^59^ and refinement in PHENIX^60,61^. The final model was assessed for quality using MOLPROBITY^62^ and figures were prepared with Pymol^63^ and ChimeraX^64^.

### Size Exclusion Chromatography-coupled Multi-Angle Light Scattering

100 μl of 2 mg/ml sample (PABC unmodified, AcK, me2K, and K606 mutants) were run on a Superdex-75 Increase 10/300 GL column (24 ml bed volume) at 0.7 ml/min in 10 mM NaH_2_PO_4_, pH 7.4; 100 mM NaF. The elution volumes varied between 12.5ml and 12.1ml, and mass was estimated for all samples between 12.6 kDa. and 15.4 kDa, informing on the monomeric status of the proteins, with no evidence of aggregation. Samples were monodispersed with Mw/Mn values of between 1.004 and 1.009.

### Circular dichroism

Circular dichroism (CD) analysis was performed on a J-810 Spectropolarimeter (JASCO) at 25°C using a path length of 1mm, over the far UV range 185–280nm. Five scans were acquired and averaged for each CD spectrum using the following parameters: data pitch, 0.1nm; bandwidth, 1nm; scan speed, 10nm/min; and response time, 2s. Solvent correction was achieved by subtraction of a buffer solution spectrum acquired with identical parameters.

### Surface Plasmon resonance (SPR)-based biosensor analysis

Direct PABC binding studies were carried out in real-time by SPR using BIACORE T200 instrument (Biacore, Uppsala, Sweden). Similar levels of His-PABC constructs were immobilized on independent channels on an NTA Sensor Chip (Cytiva) (between 200 and 400 RU coupled). A separate control flow cell was coated with Ni^2+^ and kept as a negative control to correct for refractive index changes. Binding assays were performed in a running buffer containing:1X HBSP Phosphate buffer (Cytiva), 3.4 mM EDTA and 0.005 % Tween 20. For each peptide interaction, the biosensor experiment was repeated three times. The experiments were performed at 25°C using a flow rate of 15 μl/min. Indicated peptide concentration (typical range cover at least from 20 % to 80 % saturation of the surface) were injected at 15 μl per min for 15 s. Peptide were allowed to dissociate for 10 s at 15 μl per min. After each step, surface was regenerated using 10 mM NaOH and 350 mM EDTA. Response units at equilibrium were plotted against concentration and the resulting binding curve was fitted using a 1:1 Langmuir binding model using Biacore Evaluation Software. For full-length eRF3 binding assay, the kinetic rate constants K_D_ (equilibrium dissociation constant in M), Ka (association rate constant, in M□¹ s□¹) and Kd (dissociation rate constant, in s□¹) were derived by fitting the association and dissociation phases to a simple bimolecular interaction model. Measurements were performed in triplicate.

### Cell culture and transfection

HCT116 OsTIR cells (HCT-Os) and their derivative HCT-116 PABPC1-AID (HCT-P1) cells were kindly provided by David Bartel^65^. HCT-Os and HCT-P1 cells were cultured in McCoy’s 5A Medium (Thermo) supplemented with 10 % Tet System Approved Fetal Bovine Serum (TaKaRa), 50 units/mL penicillin and 50 µg/mL streptomycin (Thermo). For degron-mediated depletion of PABPC1, HCT-P1 cells were treated overnight with 1 µg/ml doxycycline to induce OsTIR expression and then treated with 0.5 M Indole-3-acetic acid (IAA) for a minimum of 2 hours.

For expression of PABPC1 K606Q/R/A/L mutants, HCT-P1 cells were transfected with pCI-MS2V5-PABPC1 or pCI-MS2V5-PABPC1K606Q/R/A/L using Lipofectamine 3000 according to the manufacturer’s instructions for 24 hours.

### Antibody generation

Antibodies were raised in New Zealand White rabbits using K606 acetylated or dimethylated HsPABPC1-specific peptides (residues 597-608) LESPESLRS(AcK)VD or LESPESLRS(Me2K)VD, conjugated to keyhole limpet hemocyanin, with boosting using shortened SLRS(AcK)VD or SLRS(Me2K)VD peptides, as appropriate. Bleeds were checked for reactivity and antiserum from bleed 3 in each case was affinity depleted using the unmodified peptide LESPESLRSKVD. Flow-through anti-PTM antiserum was then affinity purified using either SLRS(AcK)VD or SLRS(Me2K)VD modified peptides (CovalAb).

### Immunoblotting

Cells were lysed in modified RIPA buffer [50 mM Tris-HCl (pH 7.4), 150 mM NaCl, 1 mM EDTA, 1% Igepal CA630, 0.2 % SDS, 10 mM sodium pyrophosphate, 25 mM 2-glycerophosphate, 0.5 % sodium deoxycholate, 10 mM sodium orthovanadate, 5 mM sodium fluoride, and 2 mM DTT] containing 1x Complete protease inhibitor cocktail (Roche), phosphatase inhibitor cocktail 1 (Sigma) and 1x lysine deacetylase inhibitor cocktail (MedChemExpress). Cell lysates were centrifuged at 14000 *g* for 10 min at 4°C and supernatant fractions were collected. Lysate protein concentrations were determined using BCA assay (Thermo) and equal amounts of protein were separated on Bis-Tris (4-12%) gels (Thermo) and transferred onto Immobilon-PVDF (Millipore). For immunoprecipitation analyses, equal volumes of IP eluate were used as required. Primary antibodies against eRF3a (Abcam; ab49878), V5 (Thermo; 2F11F7), AID (MBL; M214-3), His_6_-tag (Merck; MAB3834), GAPDH (Abcam; ab181602), PABPC1 (395-408, 1:1000)^66^, acK606-PABPC1, or me2K606-PABPC1 were used in PBS/0.1% Tween20 + 5% non-fat milk. Signals were detected using SuperSignal West Femto (Thermo) HRP substrate and acquired with a LI-COR Odyssey Fc Imaging System. Band intensities were quantified using Empiria Studio (versions 2.1-3.3) software. Representative blots of at least three independent experiments are shown.

### Immunoprecipitation

For co-immunoprecipitation, lysates were prepared using Immunoprecipitation Lysis buffer (ILB) [25 mM Tris/HCl (pH 8.0), 150 mM NaCl, 1 mM MgCl_2_, 0.5% Igepal CA630 and 5 mM DTT] containing 1x Complete protease inhibitor cocktail, 1x phosphatase inhibitor cocktail 1 and 1x deacetylase inhibitor cocktail for 5 min on ice. Lysates were clarified at 14000 *g* for 10 min at 4°C and supernatants retained and quantified. Equal amounts of protein in equal ILB volumes were supplemented with 5 µL RNase I_f_ (NEB) and 10 µL of a 50% slurry Protein-A Sepharose 4B beads (Cytiva) and antibody against either V5-tagged bait protein (Thermo; 2F11F7), eRF3a (Abcam; ab49878) or control rabbit IgG prior to being tumbled overnight at 4°C, washed extensively with lysis buffer followed by lysis buffer with increased salt (900 mM NaCl) at 4°C and eluted in equal volumes of 1x LDS sample buffer (Thermo) containing 2mM DTT at 75°C for 5 min.

### Confocal immunofluorescence microscopy

Cells were grown, transfected, and treated with doxycycline and/or IAA as indicated on Nunc Lab-Tek II Chamber Slide (Thermo). Cells were fixed in cold 4 % paraformaldehyde (PFA) for 10 min, permeabilised for 2 min with −20°C methanol, rehydrated in ice-cold PBS (+ 0.02% sodium azide), and treated with 3 % hydrogen peroxide for 45 min. For detection of PABPC1 and V5-tagged PABPC1, cells were blocked with 20 % normal goat serum in PBS, incubated with anti-V5 (Thermo; 2F11F7), and anti-PABPC1 (395-408) antibodies, and then incubated with AlexaFluor-546 anti-mouse (Life Technologies; A10036) and AlexaFluor-488 anti-rabbit (Life Technologies; A21206) secondary antibodies, with all washes performed in PBS. For detection of simultaneous detection of PABPC1-AID and V5-tagged PABPC1, cells were blocked with 20% normal goat serum in PBS, incubated with anti-V5 antibody, and incubated with HRP-anti-mouse secondary antibody (Pierce; #31444), with all washes performed in PBS. Opal 620 fluorophore (Akoya Biosciences) diluted in 1X Plus Amplification Diluent (Akoya Biosciences) was used to detect HRP anti-mouse signal. Thereafter, antibodies were stripped with 0.01 M sodium citrate (pH 6) at 95°C for 20 min and staining repeated using anti-AID antibody (MBL; M214-3) and Opal 540 (Akoya Biosciences). Cells were then mounted with Vectashield antifade mounting medium with DAPI (Vector Laboratories). Fluorescence images were acquired using a Zeiss LSM780 confocal laser scanning microscope. Image capture and analysis software was Zeiss Zen Black and Zen Blue v2.1.

### Statistical analysis

All statistical analyses were performed using GraphPad Prism v10 as described in text and legends.

## Acknowledgements

We thank William Richardson (deceased); Liz Blackburn, Matthew Nowicki and Martin Wear at the Edinburgh Protein Production Facility (EPPF), Coffee Xiang and David Bartel for the kind sharing of the HTC-P1 cells prior to publication, and Jason Chin for sharing the amber codon suppression/recoding plasmids. Work was funded by an MRC program (MR/J003069/1), a BBSRC project grant, (BB/P022065/1), and an MRC DTA to NKG, and a Marie Curie Fellowship 753803 to ED. AGC was supported by a Wellcome SRF (200898).

---

**Supplementary Figure 1. (A)** PaxDB quantitative mass spectrometry quantification of PABPC1 and PAM2-containing proteins in human tissues and cell lines. Violin plots depict Log(10) transformed data distribution with medians indicated by thick solid line and quartiles indicated by thin solid line. Dots represent all measurements in PaxDB as of 7/8/25. **(B)** Principal component analysis of raw PPM data used in (A).

**Supplementary Figure 2 (A)** Schematic overview of the genetic code expansion (GCE) approach using amber codon suppression for site-specific incorporation of acetyl-lysine (acK) and the protected precursor of dimethyl-lysine (BocK). **(B)** Strategy for genetically directing the site-specific incorporation of dimethyl-Lysine in recombinant PABC^33^. **(C)** Deconvoluted electrospray ionization mass spectra of PABC intermediates: (2) Boc-Cbz–protected PABC (observed mass = 14,185 Da) and (3) Cbz-protected/deprotected PABC (–100 Da) (observed mass = 14,085 Da). Expected masses: 14,185 Da (protected) and 14,085 Da (–Boc). **(D)** Deconvoluted electrospray ionization mass spectra of purified PABC variants. Observed masses: unmodified PABC, 13,994 Da; PABC-acK606 (+42 Da), 14,036 Da; PABC-me2K606 (+2×14 Da), 14,022 Da, consistent with the expected mass shifts. **(E)** Size-exclusion chromatography and SDS–PAGE analysis of purified recombinant me2K606-PABC. **(F)** SEC-MALS analysis of PABPC1 variants. Retention volumes (mL) and estimated molecular weights (Da) are indicated. **(G)** Superdex 75 size exclusion elution profiles of unmodified, acK606 and me2K606-PABC. Overlayed elution profiles are similar, indicating that the K606 status does not significantly affect the overall behaviour of this protein domain. **(H)** Circular dichroism (CD) spectra of me2K606-PABC recorded in the 190–240 nm range. The spectral shape and intensity (Δε) were compared with those of unmodified PABC (PDB ID: 3KUR), showing no significant differences in secondary structure. **(I)** Equal quantities of recombinant unmodified (un), acK606- (ac), and me2K606-PABC (me2) protein, detected using GelCode Blue protein stain (input), were immunoblotted using K606 PTM-specific antibodies.

**Supplementary Figure 3. (A)** HCT-P1 cells were either untransfected or transfected with PABPC1-V5 for 24 hours (+ doxycycline) and then either untreated or treated with IAA for 2 hours. Endogenous PABPC1-AID (green), exogenous PABPC1-V5 (red), and DNA (DAPI – blue) were detected using immunofluorescence and confocal microscopy. **(B)** HCT-P1 cells were transfected with K606X-PABPC1-V5 for 24 hours (+ doxycycline) and then treated with IAA for 2 hours. Detection as A). Individual cell zooms from the wider field are shown top-left.

**Supplementary Table 1.** Means and standard deviation data for ratiometric quantification of relative PAM2:PABPC1 levels shown in Figure 1D

**Supplementary Table 2.** acK606-PABC-eRF3a(N) crystal structure data collection and refinement statistics

