## Supplementary figures and images for "Lysine acetylation of PABPC1 C-terminal domain facilitates competitive recruitment of translation termination factor eRF3a"

### Supplemental Figure 1

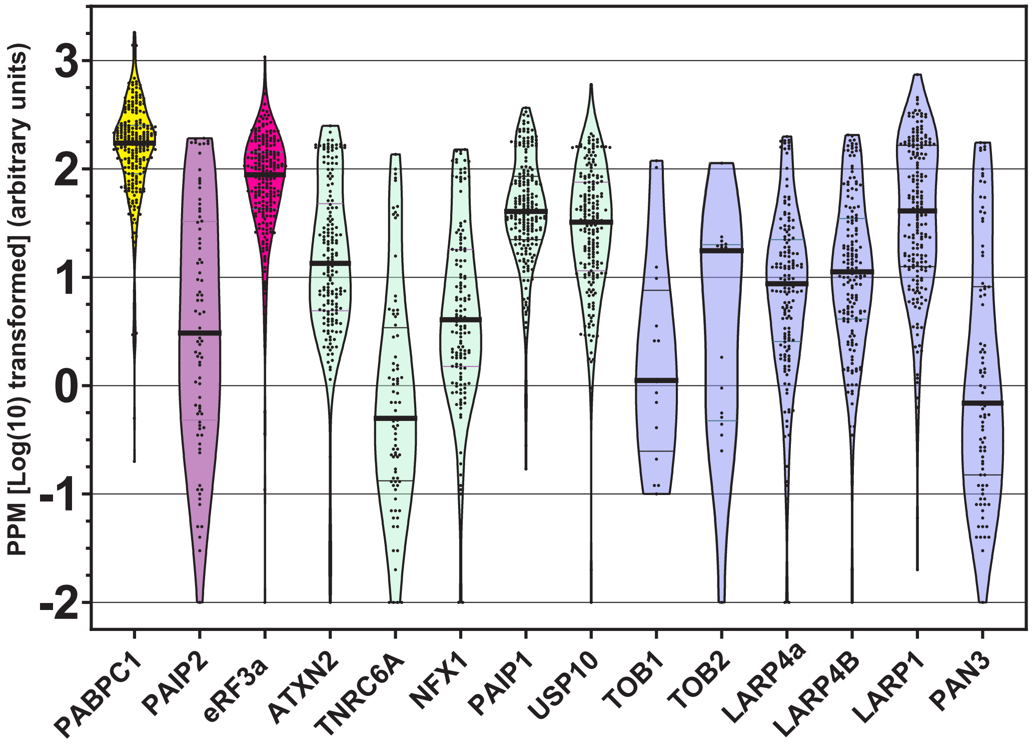

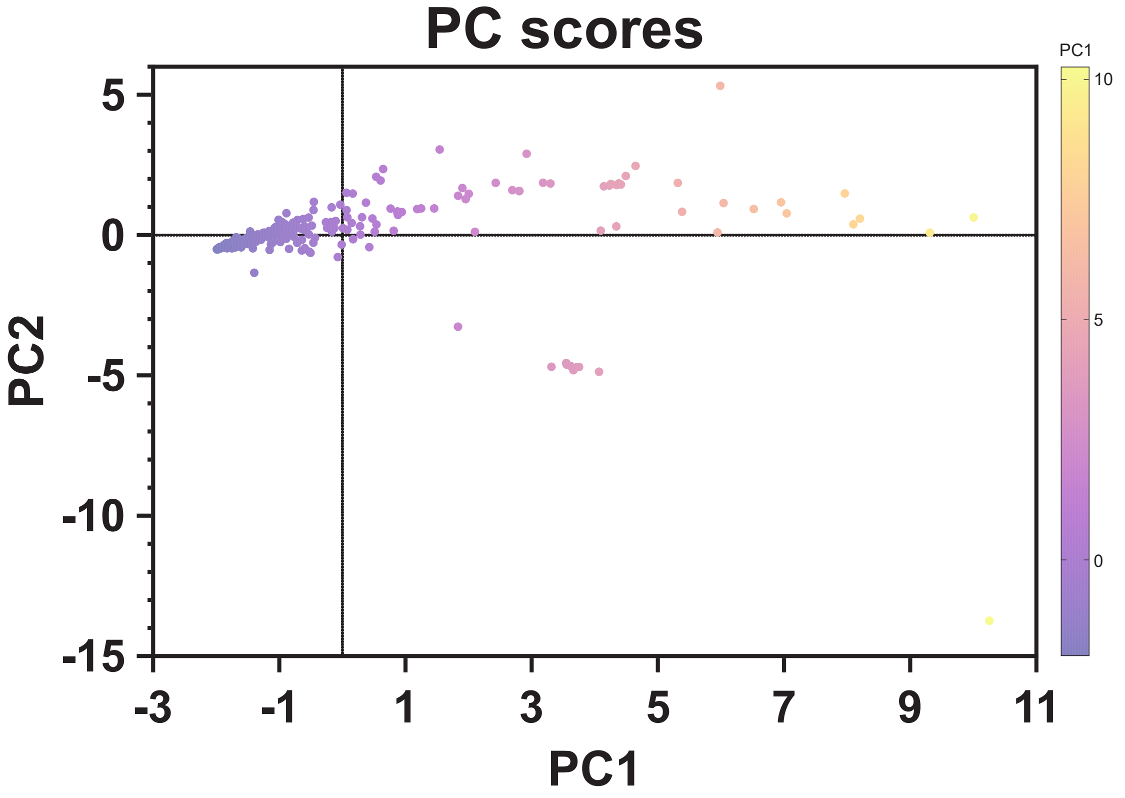

### Supplemental Figure 2

#### Supp Figure 2

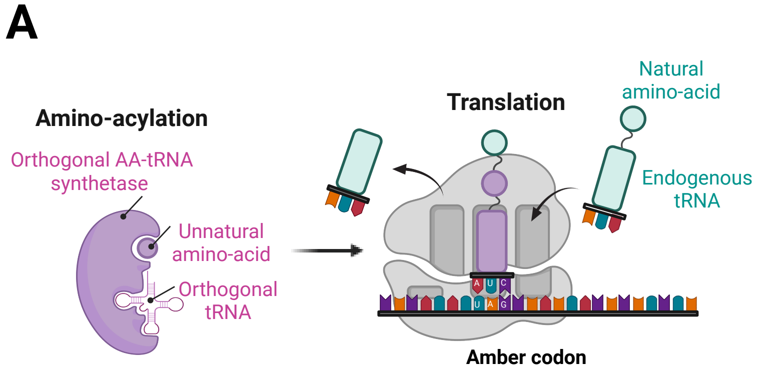


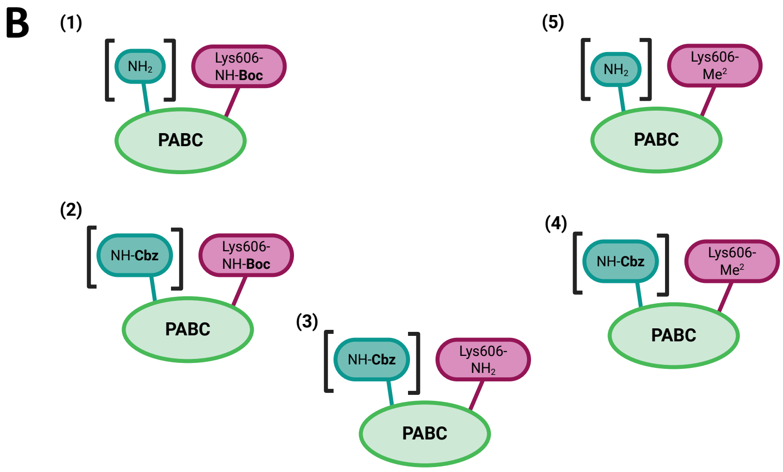


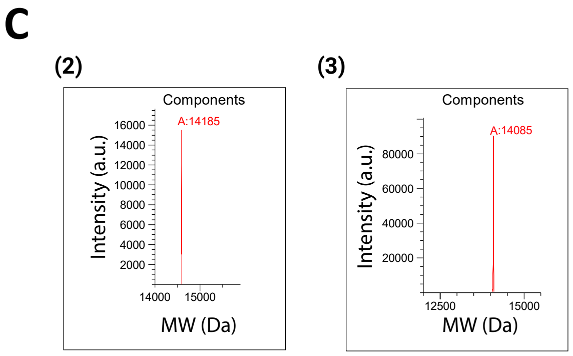


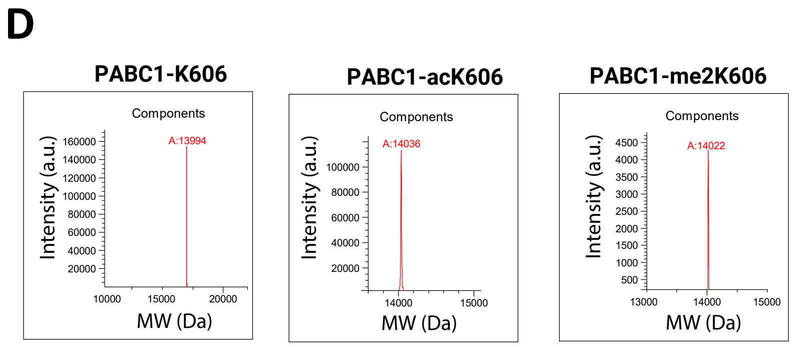


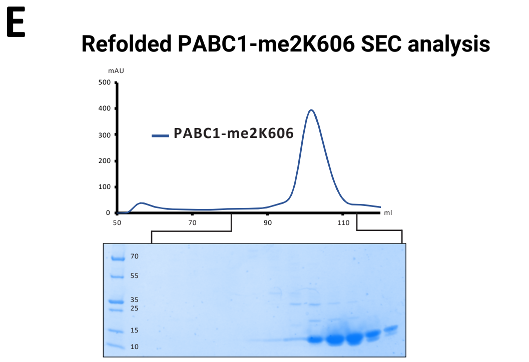


F

**SEC-MALS ANALYSIS**

| PABC1 | Ret Vol (mL) | Mw (Da) |
|---|---|---|
| UN | 13.49 | 13.98 |
| acK606 | 13.50 | 13.57 |
| me2K606 | 13.46 | 15.31 |
| Q606 | 12.88 | 13.43 |
| A606 | 12.95 | 12.99 |
| R606 | 12.8 | 12.51 |
| BSA | 10.04 | 65.65 |

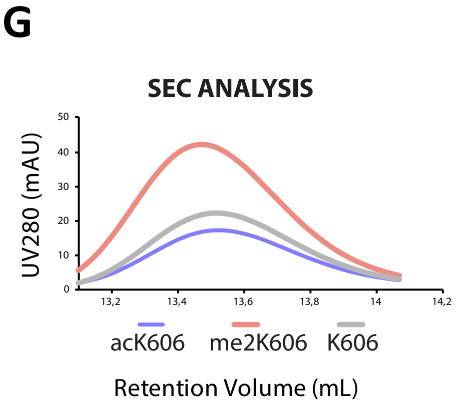


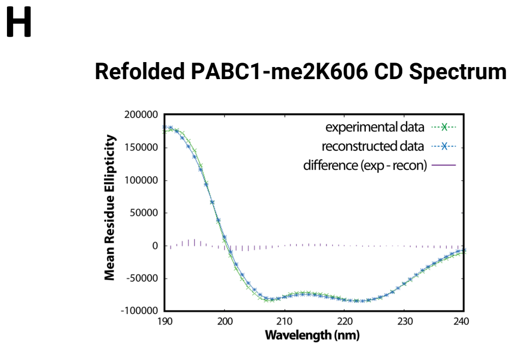


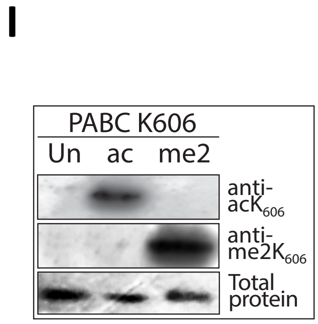

### Supplemental Figure 3

#### Supp Figure 3

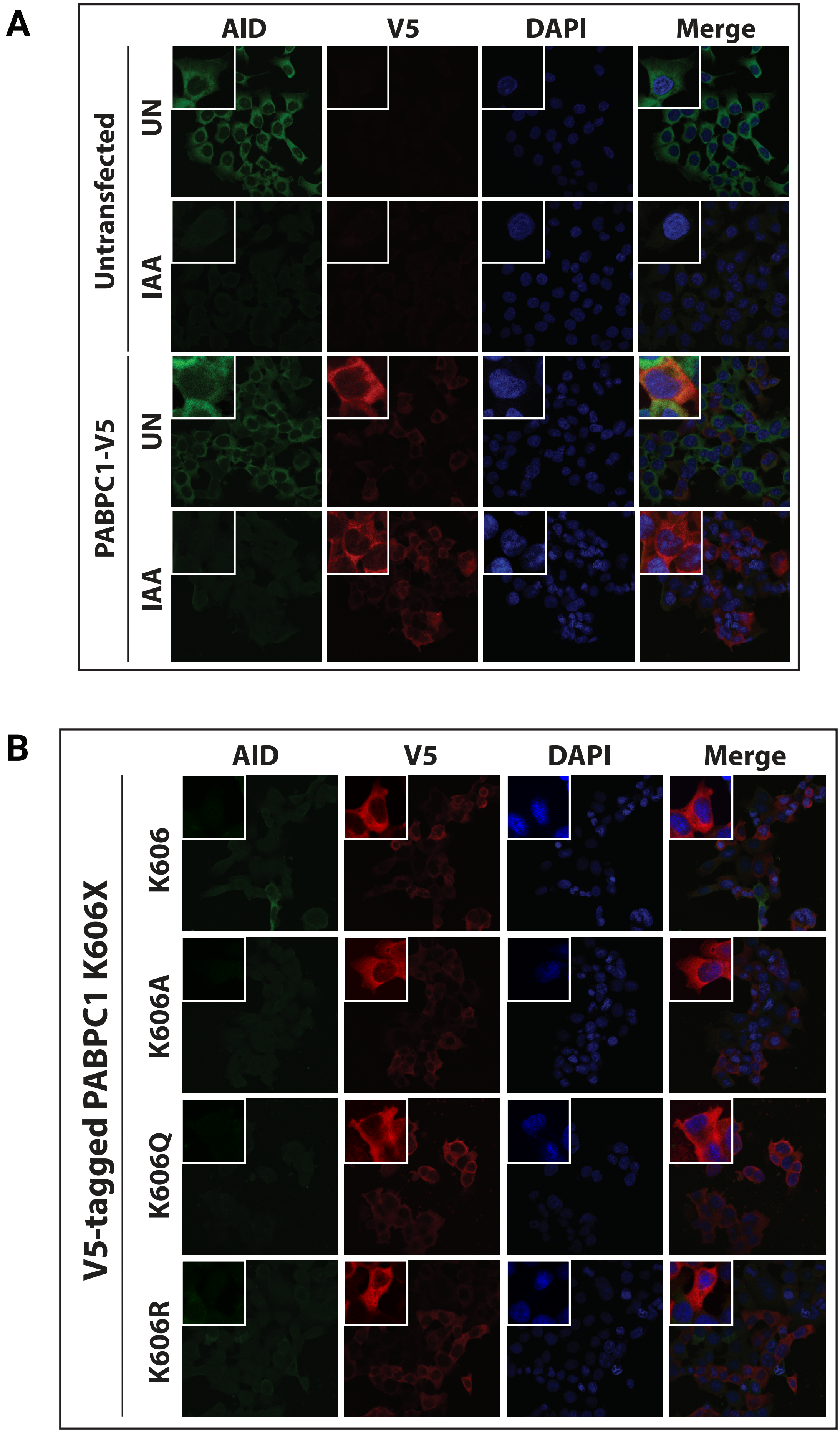
