## Supplemental Table 1 for "Lysine acetylation of PABPC1 C-terminal domain facilitates competitive recruitment of translation termination factor eRF3a"

| | PAM2:PABPC1 Mean ratio | Std. Dev |
|---|---|---|
| **PAIP2** | 0.008123 | 0.02803 |
| **eRF3a** | 0.08219 | 0.08824 |
| **ATXN2** | 0.02392 | 0.04234 |
| **TNRC6A** | 0.002677 | 0.01231 |
| **NFX1** | 0.009663 | 0.02427 |
| **PAIP1** | 0.04798 | 0.05905 |
| **USP10** | 0.03524 | 0.0509 |
| **TOB1** | 0.0008412 | 0.008127 |
| **TOB2** | 0.0009074 | 0.006466 |
| **LARP4A** | 0.01222 | 0.02957 |
| **LARP4B** | 0.01796 | 0.03509 |
| **LARP1** | 0.06032 | 0.08599 |
| **PAN3** | 0.006015 | 0.02274 |
