## Supplemental Table 2 for "Lysine acetylation of PABPC1 C-terminal domain facilitates competitive recruitment of translation termination factor eRF3a"

Supplementary Table 2: Data Collection and refinement statistics

| | |
|---|---|
| **Data Collection** | acK606-PABC1-eRF3(N) |
| Space Group | $P2\ 2_1\ 2_1$ |
| Cell Dimension | |
| a, b, c | 31.9 52.8 54.3 |
| α, β, γ (°) | 90.0 90.0 90.0 |
| Resolution (Å) | 31.89 – 1.75 |
| $R_{pim}$ | 0.058 (0.536) |
| I/σI | 7.9 (1.4) |
| Completeness (%) | 99.7 (98.8) |
| Redundancy | 6.3 (5.0) |
| $CC_{1/2}$ | 0.994 (0.719) |
| Observed refection | 64745 |
| Unique reflections | 10262 |
| **Refinement** | |
| Resolution | 31.89-1.75 |
| $R_{work}/R_{free}$ | 0.223/0.260 |
| *No atoms* | |
| Proteins | 1492 |
| Water | 48 |
| Ligands ($SO_4$) | 5 |
| *B factors (Å$^2$)* | |
| Proteins | |
| Chain A | 33.6 |
| Chain B | 41.7 |
| Ligands ($SO_4$) | 39.1 |
| Waters | 38 |
| *R.m.s Deviations* | |
| Bond lengths (Å) | 0.008 |
| Bond angles (°) | 1.505 |
| *Ramachandran plot* | |
| Favored (%) | 100 |
| Outlier (%) | 0 |
